# RecT recombinase expression enables efficient gene editing in *Enterococcus*

**DOI:** 10.1101/2020.09.01.278044

**Authors:** Victor Chen, Matthew E. Griffin, Howard C. Hang

## Abstract

*Enterococcus faecium* is a ubiquitous Gram-positive bacterium that has been recovered from the environment, food, and microbiota of mammals. Commensal strains of *E. faecium* can confer beneficial effects on host physiology and immunity, but antibiotic usage has afforded antibiotic-resistant and pathogenic isolates from livestock and humans. However, the dissection of *E. faecium* functions and mechanisms has been restricted by inefficient gene editing methods. To address these limitations, here we report the expression of *E. faecium* RecT recombinase significantly improves the efficiency of recombineering technologies in both commensal and antibiotic-resistant strains of *E. faecium* and other *Enterococcus* species such as *E. durans* and *E. hirae*. Notably, the expression of RecT in combination with clustered regularly interspaced palindromic repeat (CRISPR)-Cas9 and guide RNAs (gRNAs) enabled highly efficient scar-less single-stranded DNA recombineering to generate specific gene editing mutants in *E. faecium*. Moreover, we demonstrate that *E. faecium* RecT expression facilitated chromosomal insertions of double-stranded DNA templates encoding antibiotic selectable markers to generate gene deletion mutants. As further proof-of-principle, we use CRISPR-Cas9 mediated recombineering to knock out both sortase A genes in *E. faecium* for downstream functional characterization. The general RecT-mediated recombineering methods described here should significantly enhance genetic studies of *E. faecium* and other closely related species for functional and mechanistic studies.

**Importance:** *Enterococcus faecium* is widely recognized as an emerging public health threat with the rise of drug resistance and nosocomial infections. Nevertheless, commensal *Enterococcus* strains possess beneficial health functions in mammals to upregulate host immunity and prevent microbial infections. This functional dichotomy of *Enterococcus* species and strains highlights the need for in-depth studies to discover and characterize the genetic components underlining its diverse activities. However, current genetic engineering methods in *E. faecium* still require passive homologous recombination from plasmid DNA. This involves the successful cloning of multiple homologous fragments into a plasmid, introducing the plasmid into *E. faecium*, and screening for double-crossover events that can collectively take up to multiple weeks to perform. To alleviate these challenges, we show that RecT recombinase enables rapid and efficient integration of mutagenic DNA templates to generate substitutions, deletions, and insertions in genomic DNA of *E. faecium*. These improved recombineering methods should facilitate functional and mechanistic studies of *Enterococcus*.

## Introduction

The Gram-positive bacterial genus of *Enterococcus* has diverse origins from the environment, food, animals, and humans^1–3^. *E. faecium* has been linked to many aspects of human health, including both probiotic and pathogenic activities^1–3^. For example, *E. faecium* remains a global public health concern for its ability to acquire vancomycin resistance and cause nosocomial infections^1–3^. Yet, commensal *E. faecium* has been discovered to modulate host immunity to promote protection against other enteric pathogens, decrease diarrheal severity and has been associated with enhanced immunotherapy efficacy^4–10^. To better understand mechanisms of both pathogenic and beneficial functions of *Enterococcus*, genetic-based studies of *Enterococcus* is needed to accelerate our comprehension of these functions. Although there are some existing tools to genetically manipulate and analyze enterococci^11,12^, the study of *E. faecium* in the laboratory is largely limited by inefficient reverse genetics.

To date, the generation of scar-less genetic mutants in *Enterococcus* relies heavily on homologous recombination^11–14^. This process involves cloning homologous templates hundreds to thousands of bases long flanking the DNA edit of interest into an *Enterococcus* compatible vector that is then transformed into *Enterococcus*. Transformants are then screened for the double crossover insertion using a multistep selection and counterselection process^13^. While homologous recombination are proven ways to acquire genetic *Enterococcus* mutants, this process is laborious and takes weeks to perform. Conversely, the development of CRISPR-Cas9 gene editing methods offers new opportunities to improve scar-less mutagenesis in bacteria^15^. CRISPR-Cas9 is an endonuclease system that site-specifically cleaves genomic DNA using a homologous, complementary RNA guide^15^. The Cas9 nuclease uses guide RNA (gRNA) consisting of both CRISPR RNA (crRNA) and an antisense trans-activiating crRNA (tracrRNA) for targeted DNA cleavage^16^. In bacteria, efficient Cas9 guided genomic cleavage usually leads to cell death due to the lack of inherent DNA repair mechanisms to repair and escape Cas9 double-stranded DNA breaks. Thus, CRISPR-Cas9 can be used alongside complementary gene editing methods to select for mutants by targeting Cas9 to the wild-type, unedited genomic sequence, thereby enriching for bacteria that mutated and escaped Cas9 chromosomal cleavage. Indeed, a recent study sought to ameliorate homologous recombination selection process using CRISPR-Cas9 counterselection with in *E. faecium*^17^. While this study was successful in demonstrating CRISPR-Cas9 as an effective method for mutant selection in *E. faecium*, the method described still relied on endogenous homologous recombination, which required multiple cloning steps of both homologous template and gRNA on the same plasmid to generate a targeted mutation.

Recombineering is a gene editing method in bacteria utilizing phage-derived recombinases to mediate the incorporation of single-stranded (ss) or double-stranded (ds) DNA templates into bacterial genomes^18^. Although recombineering is well-established for model species, many bacteria including *Enterococcus* lack functional recombineering methods. More recently, recombineering has been combined with CRISPR-Cas based technologies to counterselect for successfully scar-less bacterial mutants in select species^16,19^. CRISPR-Cas mediated recombineering is advantageous over homologous recombination due to its ability to incorporate short ssDNA oligonucleotide templates that are available from commercial vendors without the need to assemble the desired template through cloning. Recombineering can also be performed with blunt end dsDNA templates for CRISPR-Cas independent recombineering. In this study, we report a recombineering system using a latent bacterial prophage RecT that is compatible with *E. faecium* and other related species (**Fig. 1**). By combining recombineering with CRISPR-Cas9 counterselection, we were able to produce both scar-less mutants via point mutations as well as controllable deletions of various sizes. Additionally, we report our system to have gene editing activity with PCR generated dsDNA templates. Overall, we show the versatility of our RecT-mediated recombineering method to produce substitution, deletion, and insertion mutants in *E. faecium* to enable facile genetic-based studies.

**Figure 1.**
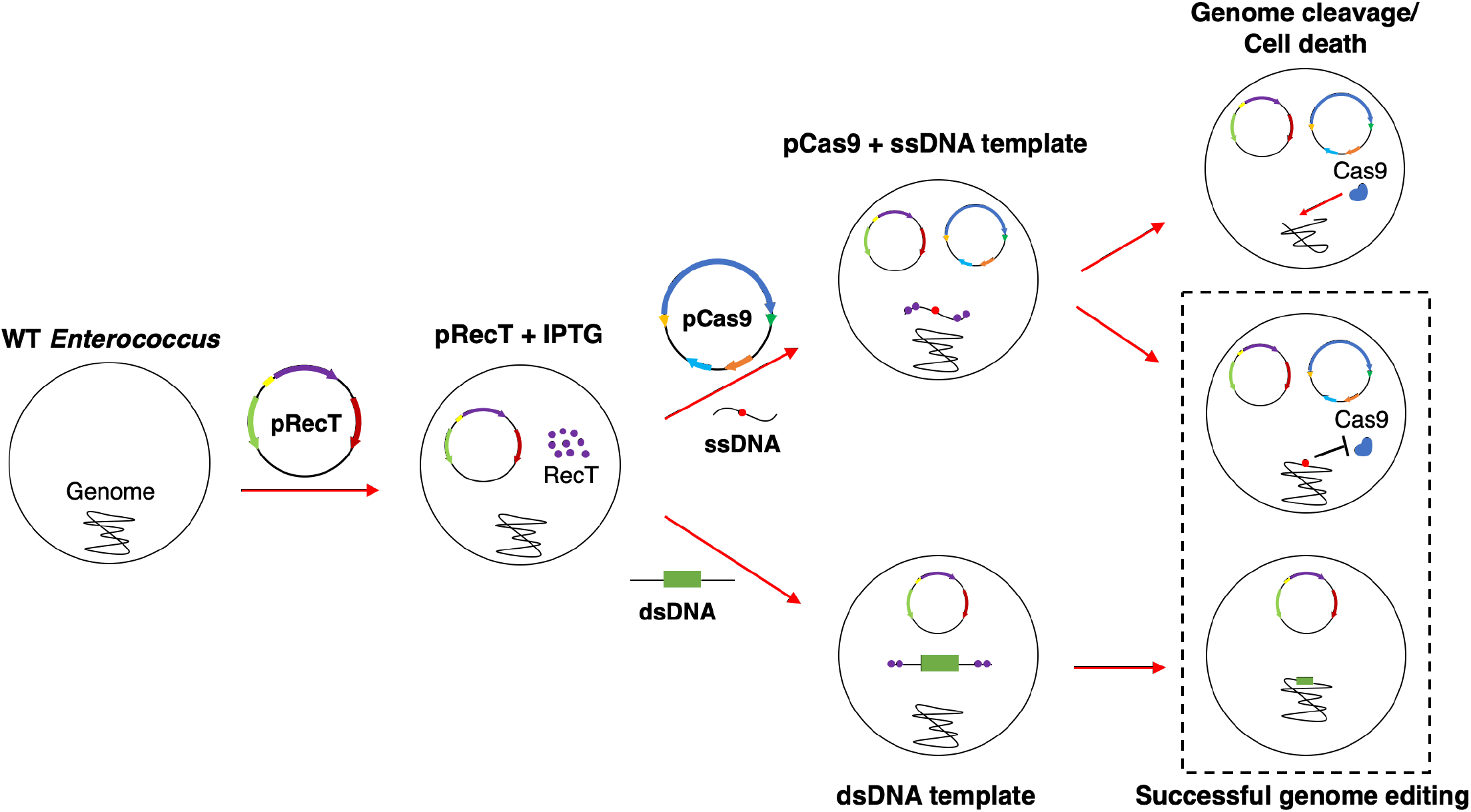
Scheme of recombineering in *Enterococcus* using CRISPR-Cas9 counterselection or genome inserted antibiotic selection. pRecT or pRecT_2 harboring cells were induced for RecT production via IPTG. RecT producing cells were transformed with either ssDNA template and pCas9 encoding genome targeting gRNA, or dsDNA template encoding an antibiotic selection marker flanked by homology arms targeting the genome. Cells that successfully recombined with template escape Cas9 dsDNA cleavage or were antibiotic resistant mediated by genomically inserted antibiotic cassette. Cells that do not recombine had their genomes cleaved by Cas9, which leads to cell death or were antibiotic sensitive and do not grow.

## Results

### S. pyogenes Cas9 nuclease is functional in E. faecium

To use CRISPR-Cas as a selection tool for *Enterococcus* genetic engineering, we first tested whether the Cas9 nuclease protein could be functionally expressed in the commensal *E. faecium* strain Com15. We based our expression system on the pVPL3004 vector, which has previously been used for CRISPR-Cas9 mediated recombineering in *Lactobacillus reuteri*^19^. The vector pVPL3004 encodes *S. pyogenes cas9*, erythromycin resistance cassette *ermC, tracrRNA*, and a crRNA cloning site. To facilitate cloning, we added a high copy number origin of replication from pUC19 into pVPL3004, which allowed for successful amplification and cloning in *E. coli*. Additionally, we found that erythromycin selection for pVPL3004 to be unreliable, as spontaneously resistant colonies formed over a few days after transformation of this plasmid. Thus, we exchanged the erythromycin resistance cassette for the chloramphenicol resistance gene, chloramphenicol acetyltransferase (*cat*). The resulting plasmid pCas9 encodes for *S. pyogenes Cas9, tracrRNA, crRNA, cat*, and the pUC19 origin or replication (**Fig. 2A**). pCas9 was transformed into *E. faecium* via electroporation of lysozyme treated cells, and transformants were assayed for Cas9 expression. Western blot analysis confirmed that *E. faecium* harboring pCas9 expresses Cas9 protein as opposed to the wild-type control (**Fig. 2B**). Furthermore, other *Enterococcus* species transformed with pCas9 also successfully expressed Cas9, indicating that this vector can be used to deliver CRISPR-Cas9 machinery across other *Enterococcus* species (**Fig. S1**).

**Figure 2.**
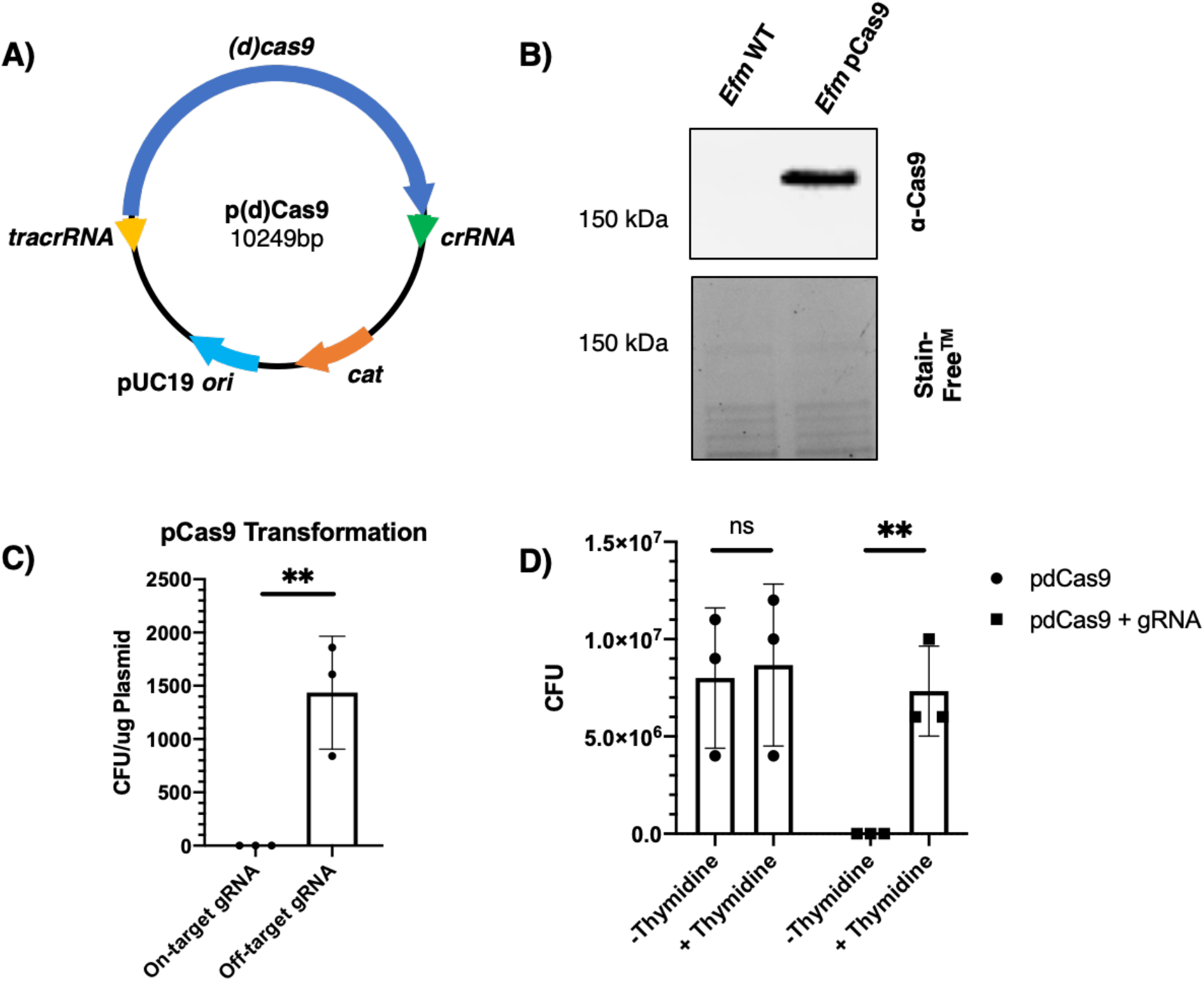
Cas9 expression and activity characterization in *E. faecium*. A) pCas9 and pdCas9 plasmid map is shown. Plasmids encode *S. pyogenes cas9, tracrRNA, crRNA*, pUC19 origin of replication and chloramphenicol acetyltransferase (*cat*). B) Western blot analysis was performed on *E. faecium* Com15 harboring pCas9, using wild-type (WT) as a control. HRP-conjugated anti-Cas9 was used (Cell Signaling Technologies). Corresponding Stain-Free^™^ gel is shown. Single band appears above 150 kDa. C) Transformation of pCas9 encoding on-target or off-target gRNA was transformed into *E. faecium*. CFU was counted and normalized for the amount of plasmid added. On-target gRNA targets *thyA*, while off-target gRNA is nearly identical to that of on-target’s but with two bases mutated at the 3’ end. D) *E. faecium* harboring pdCas9 encoded with or without gRNA targeting *thyA* was grown on MM9YEG agar with or lacking thymidine. CFU counts represent the number cells in 5 μL of culture.

Previous studies have indicated that Cas9 targeting of genomic DNA is lethal in bacteria^15^. Additionally, Cas9 nuclease activity is highly sensitive to mismatches between the gRNA and the 3’ region of the targeted sequence^16^. To confirm that Cas9 is active in *E. faecium*, we performed transformation assays of pCas9 into *E. faecium* with a gRNA that targeted thymidylate synthase (*thyA*) (pCas9-*thyA*). As a control, we created a separate pCas9 vector encoding an off-target gRNA by mutating two nucleotides at the 3’ end of the on-target gRNA. As expected, transformation with pCas9-*thyA* into *E. faecium* yielded very few colonies (**Fig. 2C**). In contrast, transforming pCas9 encoding the off-target gRNA yielded significantly more colonies, indicating Cas9 is active in *E. faecium* and kills the bacteria post-transformation (**Fig. 2C**). We also examined whether catalytically dead Cas9 (dCas9) could also be used for gene knockdown by CRISPR interference (CRISPRi)^20^. Mutation of the *Cas9* active sites, D10A and H840A, yielded and incorporation of the *thyA* targeting gRNA yielded pdCas9-*thyA*, which was then transformed into *E. faecium*. Resulting transformants were then grown in thymidine-rich liquid BHI medium and plated onto MM9YEG agar medium with or without thymidine. Colonies from the resulting plates show *E. faecium* was able to grow on medium containing thymidine but not minimal medium alone, indicating dCas9 actively repressed the expression of thymidylate synthase to cause thymidine auxotrophy (**Fig. 2D**).

### RecT expression enables efficient CRISPR-Cas9 mediated recombineering in *E. faecium* Com15

Recombineering methods in various bacteria utilize RecT proteins to incorporate introduced DNA templates^21^. In nature, RecT proteins are ssDNA annealing proteins that function in bacteriophage recombination^22^. Because RecT proteins can exhibit species-specific activity, we reasoned that prophage operons in *E. faecium* may be a useful source for finding RecT proteins that are active and compatible with enterococci. Within the *E. faecium* Com15 genome, we found one *recT* gene (EFWG_RS08525) in a prophage operon. To ectopically express RecT for recombineering, we cloned *recT* to be under the control of an IPTG inducible promoter in a plasmid (**Fig. 3A**). The plasmid pRecT also contains the genes *lacI* and *ermC* for IPTG transcriptional control and erythromycin selection, respectively (**Fig. 3A**). With both a CRISPR-Cas9 delivery vector and a RecT expression vector in hand, we next investigated whether we could genetically manipulate *E. faecium* Com15 using CRISPR-Cas9 mediated recombineering. We first transformed *E. faecium* Com15 with pRecT and selected for transformants using erythromycin on BHI agar. *E. faecium* containing pRecT were then grown in liquid culture and induced with IPTG for ectopic RecT expression until mid-log phase, before being harvested and co-transformed with ssDNA template and pCas9-*thyA*. Once transformed, cells were selected using chloramphenicol on BHI agar as a proxy for Cas9 expression and screened for recombination events (**Fig. 1**).

**Figure 3.**
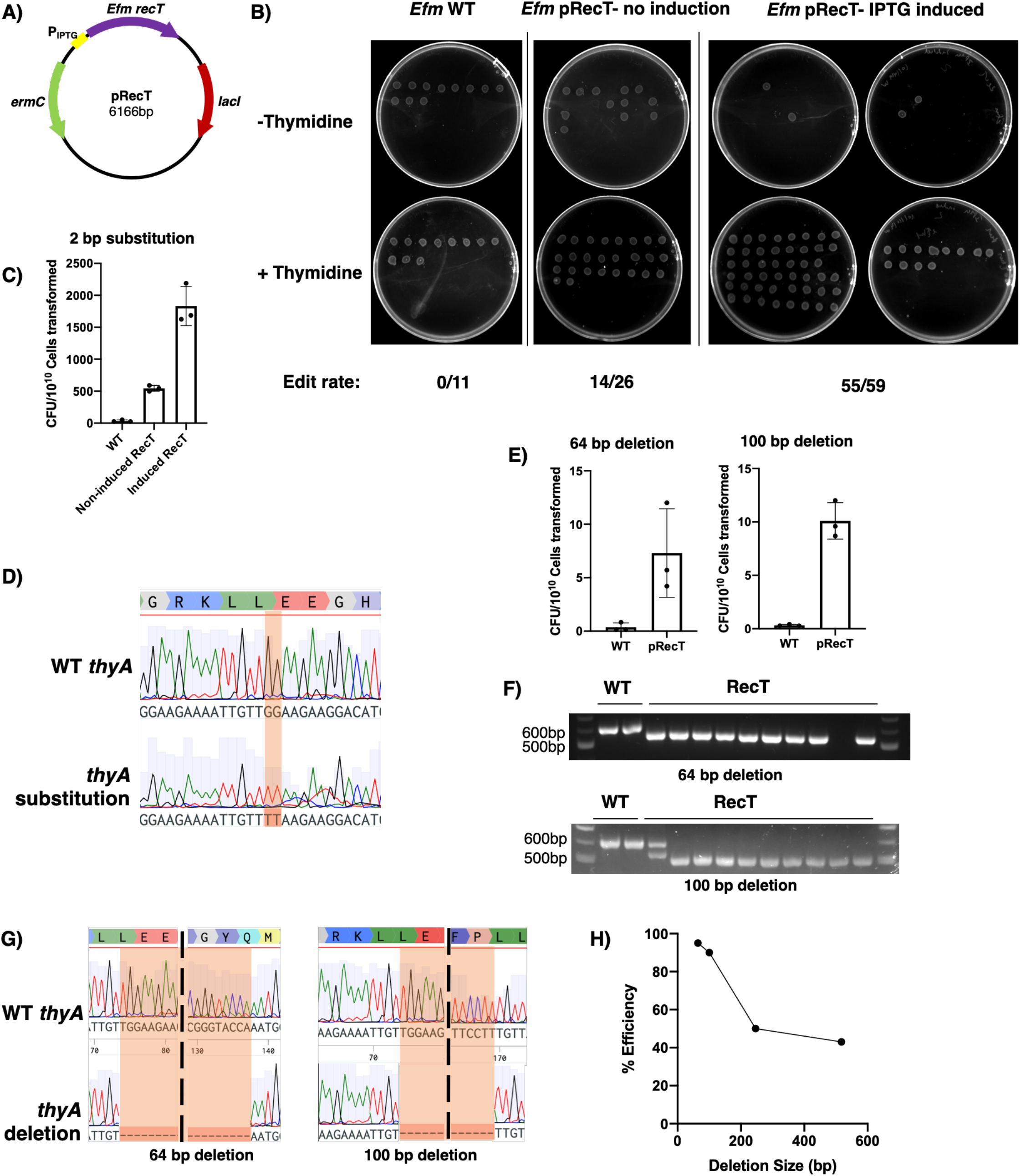
*E. faecium* Com15 CRISPR-Cas9 mediated recombineering. A) Plasmid map of pRecT is depicted. pRecT encodes for *recT* derived from the genome of *E. faecium* under an IPTG inducible promoter, *ermC* and *lacI*. B) Spot assay of resulting CRISPR-Cas9 mediated recombineering experiment to induce two base substitution in *E. faecium* Com15 (*Efm*) *thyA* is depicted. *E. faecium* Δ*thyA* mutants are unable to grow on MM9YEG media lacking thymidine. C) Relative CFU for two base pair substitution experiment is shown. CFU is normalized to 10^10^ *E. faecium* transformed in the experiment. D) Sequence identity of two base pair substitution matches with the proposed induced mutation. E) Relative CFU per 10^10^ *E. faecium* cells transformed is shown for *thyA* 64bp and 100bp deletion mutants made via CRISPR-Cas9 mediated recombineering. F) DNA gel of colony PCR of resulting mutants from E). Mutants show a modest gel shift corresponding to the deletion size on DNA gel. G) Sequence identity of deletion mutants is shown to be of high fidelity, with the exact number of proposed nucleotides deleted for both 64bp and 100bp deletions. H) Editing efficiency was measured against size of deletion. %Efficiency was calculated via correctly recombineered colonies/total number of colonies x100%. Sizes of deletions depicted are 64bp, 100bp, 256bp and 517bp.

To test for site-specific mutagenesis, we first designed ssDNA oligonucleotides that introduced two base-pair substitutions into *thyA* that would produce an early stop codon and ablate the protospacer-adjacent motif (PAM) sequence to prevent Cas9 cleavage^16^. To determine whether RecT improved recombineering, we performed the experiment using wild-type *E. faecium* and *E. faecium* harboring pRecT grown with or without IPTG induction. Colonies that grew after transformation with pCas9-*thyA* were screened by spotting on MM9YEG agar with or without thymidine. We found that in the wild-type condition, none of the colonies selected were thymidine auxotrophs. Conversely, RecT-expressing *E. faecium* successfully produced thymidine auxotrophs, with the non-induced and IPTG induced conditions yielding 53.8% and 93% editing efficiencies, respectively (**Fig. 3B**). These data suggest that the ectopic expression of RecT greatly improves recombineering activity in *E. faecium*. Consistently, we found that the number of colony formation units as a result of the recombineering transformation for *E. faecium* was higher in the IPTG induced condition as opposed to the uninduced or wild-type conditions (**Fig. 3C**). We confirmed that the mutant genotype sequences contained the desired two base-pair substitutions in *thyA*, indicating that this system is both efficient and accurate in producing substitution mutations in *E. faecium* (**Fig. 3D**).

Although we successfully generated substitutions via recombineering, our experiments show a low, basal level of spontaneous CRISPR-Cas9 escape that may make screening for other genes without obvious phenotypes more difficult. To address this concern, we asked whether PCR amplification was a comparable method to screen for recombineering. Here, we designed oligonucleotide templates that would generate short deletions (64 and 100 bp) in *E. faecium thyA* that were discernable by agarose gel analysis after PCR amplification. Using these new templates with pCas9-*thyA* and pRecT, we found that colony formation units were significantly higher in RecT-expressing cells compared to wild-type cells (**Fig. 3E**). Using spot assays as described above, we found that the rates of on-target recombineering were 95% and 90% for the 64 and 100 bp deletions, respectively (**Fig. S2A**). We found that wild-type *E. faecium* also produced thymidine mutants without the help of RecT at a markedly lower rate of efficiency, a phenomenon that has been previously described in *Staphylococcus aureus*^21^ (**Fig. S2A**). Colonies were also screened via colony PCR for short deletions. Compared to the wild-type cells, PCR amplification of the recombineered cells showed observable gel shifts corresponding to the proper deletion sizes (**Fig. 3F**). Sanger sequencing of resulting thymidine auxotrophs show deletions in *thyA* of expected size in both 64 and 100 bp deletions, indicating the generation of deletions via CRISPR-Cas9 mediated recombineering is of high fidelity (**Fig. 3G**). We further performed additional gene deletion experiments to approximate the maximum deletion size achievable using 100 base ssDNA oligonucleotide templates in CRISPR-Cas9 mediated recombineering. We found that this technology is able to generate deletions of up to at least 517bp with reduced editing efficiency (**Fig. 3H**). To determine if this editing system can be applied to closely related species, we applied our CRISPR-Cas9 mediated recombineering protocol to *Enterococcus durans, Enterococcus hirae* and *Enterococcus mundtii*. We were able to generate small deletions in their respective *thyA* genes for *E. durans* and *E. hirae* using CRISPR-Cas9 mediated recombineering without significant modification or optimization of our standardized approach (**Fig. S2B-G**). This indicates our developed technology can be readily applied to other *Enterococcus* species as well.

### RecT recombineering enables the generation of insertion mutants in *E. faecium* Com15

To test if our CRISPR-Cas9 mediated recombineering system is able to produce insertion mutants, we designed commercially synthesized ssDNA oligonucleotide templates that contained small DNA insertions of 15 bases into the *E. faecium* genome. However, any attempts to produce insertions with the commercial ssDNA templates proved unsuccessful, likely because ssDNA oligonucleotide templates are too short to effectively make insertions. Thus, we attempted to make insertions using longer dsDNA templates (**Fig. 1**). We cloned a 787 bp chloramphenicol acetyltransferase (*cat*) operon flanked by two 1kb homology arms homologous to *E. faecium thyA* into the plasmid pET21 (**Fig. 4A**). To generate long DNA template, we PCR amplified the cloned DNA template and purified the DNA products for transformation into *E. faecium* Com15. To avoid complications due to chloramphenicol resistance from pCas9-*thyA*, we performed the recombineering experiment without the help of Cas9 selection, relying solely on the *cat* operon insertion for chloramphenicol-based selection. Transformed, chloramphenicol-resistant colonies were spot assayed as described above. We found that 94% of the tested colonies showed thymidine auxotrophy with only one chloramphenicol-resistant colony that was unable to grow on chloramphenicol containing MM9YEG plates, suggesting nearly complete on-target insertion of *cat* into the *thyA* locus (**Fig. 4B**). Although previous studies have indicated that dsDNA templates required the exonuclease RecE in combination with RecT to produce dsDNA recombineering mutants in bacteria^18,23^, our results suggest that insertional mutants could be produced in our system with only dsDNA template and RecT. Notably, no colonies were formed in cells lacking pRecT, indicating that insertional mutations using this method are highly dependent on RecT expression. Additionally, PCR of *thyA* in mutant colonies showed a gel shift of about 787bp, which corresponded to the desired *cat* insertion size (**Fig. 4C**). Sanger sequencing further shows that the *cat* insert into *thyA* in *E. faecium* was placed in the expected region at high fidelity (**Fig. 4D**).

**Figure 4.**
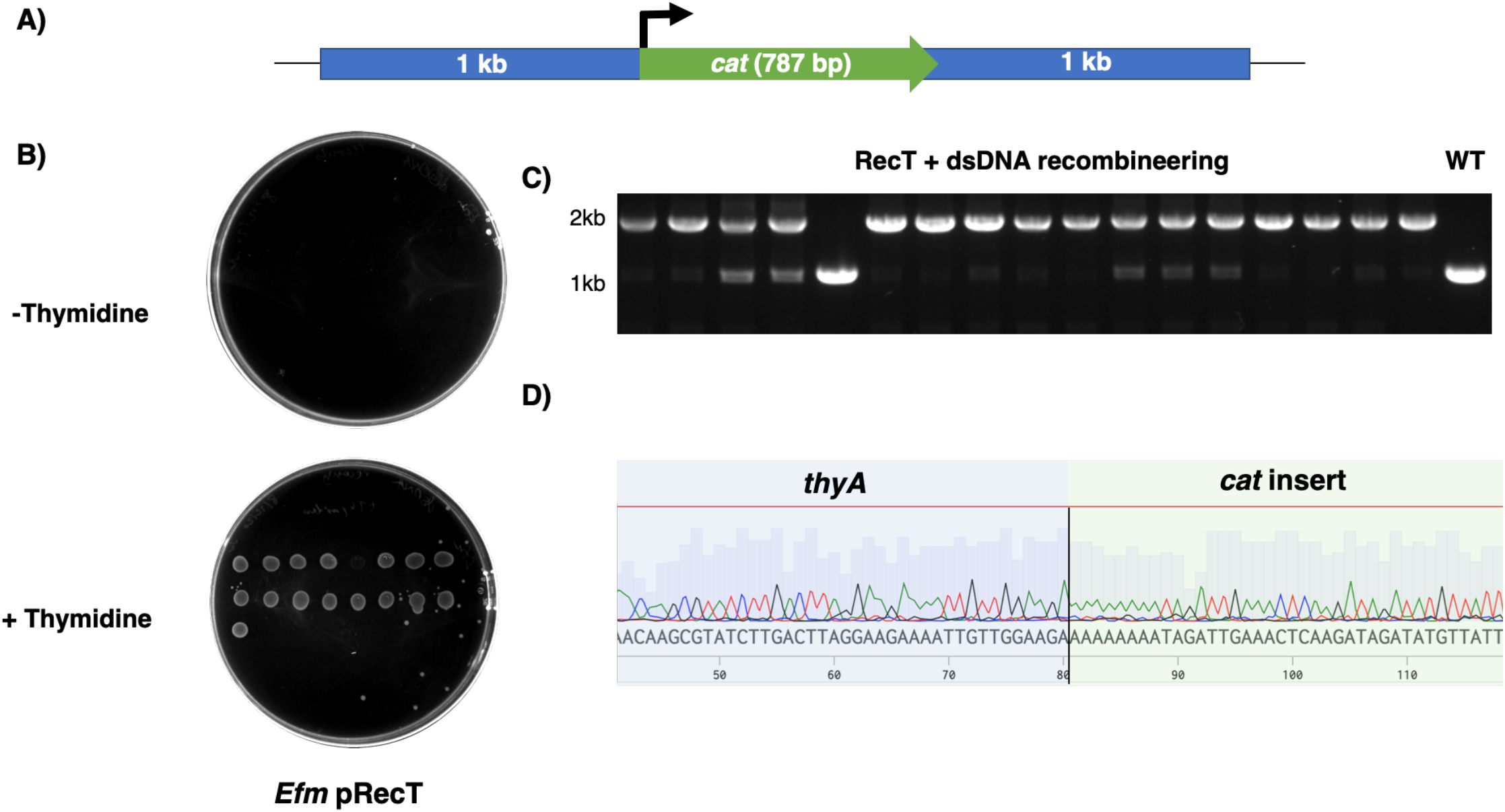
Recombineering of long DNA template enables the insertion mutations. A) Scheme of template depicts chloramphenicol acetyltransferase under the control of the constitutive *bacA* promoter, flanked by 1kb homology arms targeting *thyA* in *E. faecium*. Fragment was cloned into pET21 for PCR amplification. *bacA* promoter was used for constitutive expression of *cat*. B) Spot assay of colonies resulting from long dsDNA recombineering is depicted from *E. faecium* Com15 harboring pRecT. C) DNA gel of colony PCR of resulting colonies produced in dsDNA recombineering. WT *E. faecium* is included as a control. Notably, one colony picked exhibited a band corresponding to WT, but is unable to grow on screening plates because of the lack of inserted *cat* gene. D) Sequence identity of mutants show *cat* insertion in *thyA* at the correct location for dsDNA recombineering.

### CRISPR-Cas9 mediated recombineering can be used to efficiently generate double knockouts

To demonstrate the utility of our CRISPR-Cas9 mediated recombineering method in efficiently generating multi-gene knockouts, we chose sortase A (*srtA*) as our gene target of interest. Importantly, Sortase A is a protein that is responsible for attaching many cell surface proteins to the cell walls of Gram-positive bacteria^24^. In enterococci, sortase-dependent proteins have been found to be responsible for adhesion, biofilm formation and host colonization^25–27^. In our genome analyses, we found that *E. faecium* Com15 encodes two copies of *srtA* that we named *srtA-1* (EFWG_RS17355) and *srtA-2* (EFWG_RS05700) for this study (**Fig. 5A**). Within the genome, *srtA-1* is localized near genes that encode proteins that do not appear to be cell wall associated (**Fig. 5A**). However, a pilin formation operon can be found next to *srtA-2* (**Fig. 5A**). Pili have been previously shown to be important in Gram-positive bacteria for pili formation and is dependent of *srtA* for attachment on the cell wall^28,29^. To test the effects of knocking out *srtA* in *E. faecium*, we generated single and double knockouts of *srtA-1* and *srtA-2* in *E. faecium* Com15 using our CRISPR-Cas9 mediated recombineering method as previously described (**Fig. 5B and 5C**). To generate knockouts, we produced a 76bp deletion in *srtA-1* (**Fig. 5B**) and a 62bp deletion in *srtA-2* (**Fig. 5C**), with the double knockout strain harboring both mutations (**Fig. 5B and 5C**). In order to generate the Δ*srtA-1/2* double knockout strain, we took the Δ*srtA-1* mutant and cured its CRISPR and recombineering plasmids. In brief, cells were passaged, streaked for single colonies, and screened for erythromycin and chloramphenicol sensitivity over the course of three days. Cured *E. faecium* Δ*srtA-1* cells were then re-transformed with pRecT before knocking out *srtA-2* using CRISPR-Cas9 mediated recombineering. To assess the phenotypic effects of these knockouts, we performed light microscopy and observed that the knockouts had modest cell chaining effects when compared to the wild-type condition (**Fig. 5C**). To investigate whether the cell chaining effects were due to *srtA* knockouts, we performed quantitative analyses of the cell chains as previously described^30^ using isogenic knockout strains and control. To produce true isogenic mutants, we cured all the plasmids (pRecT and pCas9) from our knockout strains, using a pRecT cured *E. faecium* Com15 strain as our wild-type background control. Notably, we found that all of the mutants produced some level of increased chaining when compared to the control (**Fig. 5D and S3A**). However, the Δ*srtA-1/2* double knockout strain yielded an increased chaining effect when compared to both Δ*srtA-1 and* Δ*srtA-2* strains (**Fig. 5D**). To reinforce these results, we created complementation plasmids by cloning either *srtA-1* or *srtA-2* constitutively expressed under the *bacA* promoter into pKH12^14^ to generate the resulting plasmids psrtA1 and psrtA2 (**Fig. S3B**). psrtA1 and psrtA2 were then transformed into *E*. faecium ΔsrtA*-1/2* and assayed for chaining. Overall, we found the complementation strains formed shorter chains than the *E. faecium* ΔsrtA*-1/2* double knockout strain (**Fig. 5D**), with chaining levels comparable to that of *E. faecium* ΔsrtA*-1* and *E. faecium* ΔsrtA*-2* (**Fig. S3C**). Overall, these proof-of-principle experiments highlight the versatility of our approach to rapidly assay gene function in cells.

**Figure 5.**
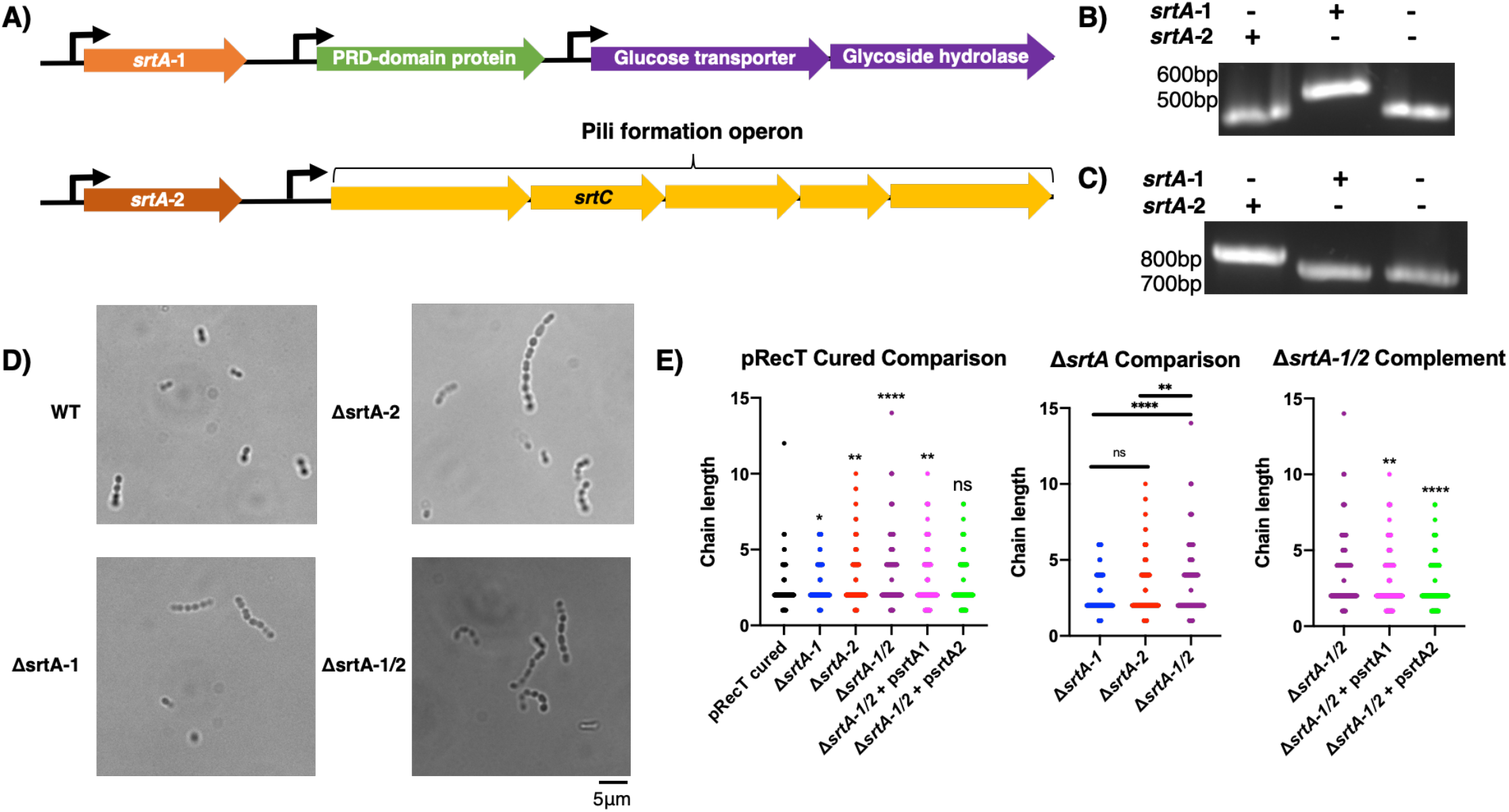
CRISPR-Cas9 mediated recombineering enables the creation of double knockouts in *E. faecium*. A) *E. faecium* Com15 encodes two copies of sortase A in separate parts of the genome. *srtA-1* localizes around genes that are seemingly unrelated to cell surface proteins. *srtA-2* localizes next to a pilin formation operon that encodes a copy of pilin forming *srtC*. B) DNA gel showing PCR of *srtA-1* on *E. faecium* Δ*srtA-1*, Δ*srtA-2* and Δ*srtA-1/2* mutants. Gel shifts corresponding to a 76bp deletion can be observed in Δ*srtA-1* and Δ*srtA-1/2* mutants when compared to Δ*srtA-2* that has the WT form of *srtA-1*. C) DNA gel showing PCR of *srtA-2* on *E. faecium* Δ*srtA-1*, Δ*srtA-2* and Δ*srtA-1/2* mutants. Gel shifts corresponding to a 62bp deletion can be observed in Δ*srtA-2* and Δ*srtA-1/2* mutants when compared to Δ*srtA-1* that has the WT form of *srtA-2*. D) Light microscopy representing *E. faecium* Com15 WT, Δ*srtA-1*, Δ*srtA-2* and Δ*srtA-1/2* mutants. Increased chaining effects can be seen in the mutant strains when compared to the WT. E) Quantification of chain length for control, mutants and complement was performed. pRecT cured *E. faecium* Com15 was used for the control condition. Approximately 300 particles were picked for each condition. Kruskal-Wallis ANOVA with Dunn’s correction was used to statistically compare multiple conditions. Graphs showing chaining comparisons for pRecT vs all other conditions, *E. faecium* Δ*srtA*, and *E. faecium* Δ*srtA-1/2* vs complement are shown. Further comparisons are graphed in Figure S3C.

### Recombineering can be applied to clinically relevant vancomycin-resistant enterococci (VRE)

Previous literature has suggested that human associated *E. faecium* are phylogenetically separated into two distinct clades, Clade A1 and Clade B, representing pathogenic and commensal *E. faecium*, respectively^31,32^. To determine whether our recombineering technology is compatible with clinically relevant, multi-drug resistant, Clade A1 *E. faecium*, we chose two vancomycin-resistant *E. faecium* strains, *E. faecium* TX0082 and *E. faecium* ERV165. Unfortunately, we found that vancomycin-resistant *E. faecium* are commonly naturally resistant to erythromycin^33^, rendering them incompatible with our original pRecT plasmid. To alleviate this, we transferred the IPTG inducible *recT* operon from pRecT onto the vector plZ12, which harbors a spectinomycin resistance gene, to produce pRecT_2 (**Fig. S4A**). Given our previous observations of highly variable transformation efficiencies across different *Enterococcus* species and strains, we also chose to further optimize our transformation protocol by applying either our original protocol of lysozyme-mediated cell wall degradation or overnight growth in high concentrations of glycine.^34^

Recombineering of the two VRE strains was performed using a purified PCR product containing a *cat* operon flanked by 1kb homology arms targeting *thyA*. Resulting colonies from both transformation methods were counted and screened for the *cat* insert in their *thyA* genes using PCR. In both *E. faecium* TX008 and ERV165 strains, we found that cells harboring pRecT_2 produced more transformants per cells transformed than did their respective wild-type controls (**Fig. 6A and 6B**). In general, we found the glycine transformation method outperformed the recombineering efficiency of the lysozyme transformation method in both strains (**Fig. 6A and 6B**). Surprisingly, we observed several colonies formed in the wild-type condition without pRecT_2 using the glycine transformation method but not in the lysozyme transformation method in both VRE strains (**Fig. 6A and 6B**). To check if the correct insertion was made, we picked colonies formed by all conditions and performed PCR analysis on the *thyA* region in the genome. We found that for *E. faecium* TX0082, every colony possessed an insertional mutation corresponding to the size of the *cat* operon (**Fig. 6C and S4B**). Similarly, nearly all colonies of *E. faecium* ERV165 possessed an insertional mutation corresponding to the size of the *cat* operon (**Fig. 6C and S4C**). To validate the insertional mutations, we performed Sanger sequencing on a subset of the PCR products produced in each condition and found that every sampled amplicon possessed the *cat* operon insertion in the expected region of the *thyA* gene (**Fig. 6D**). We found that the wild-type VRE strains were unable to grow effectively on MM9YEG screening plates, which precluded functional validation of *thyA* auxotrophy by spot assays.

**Figure 6.**
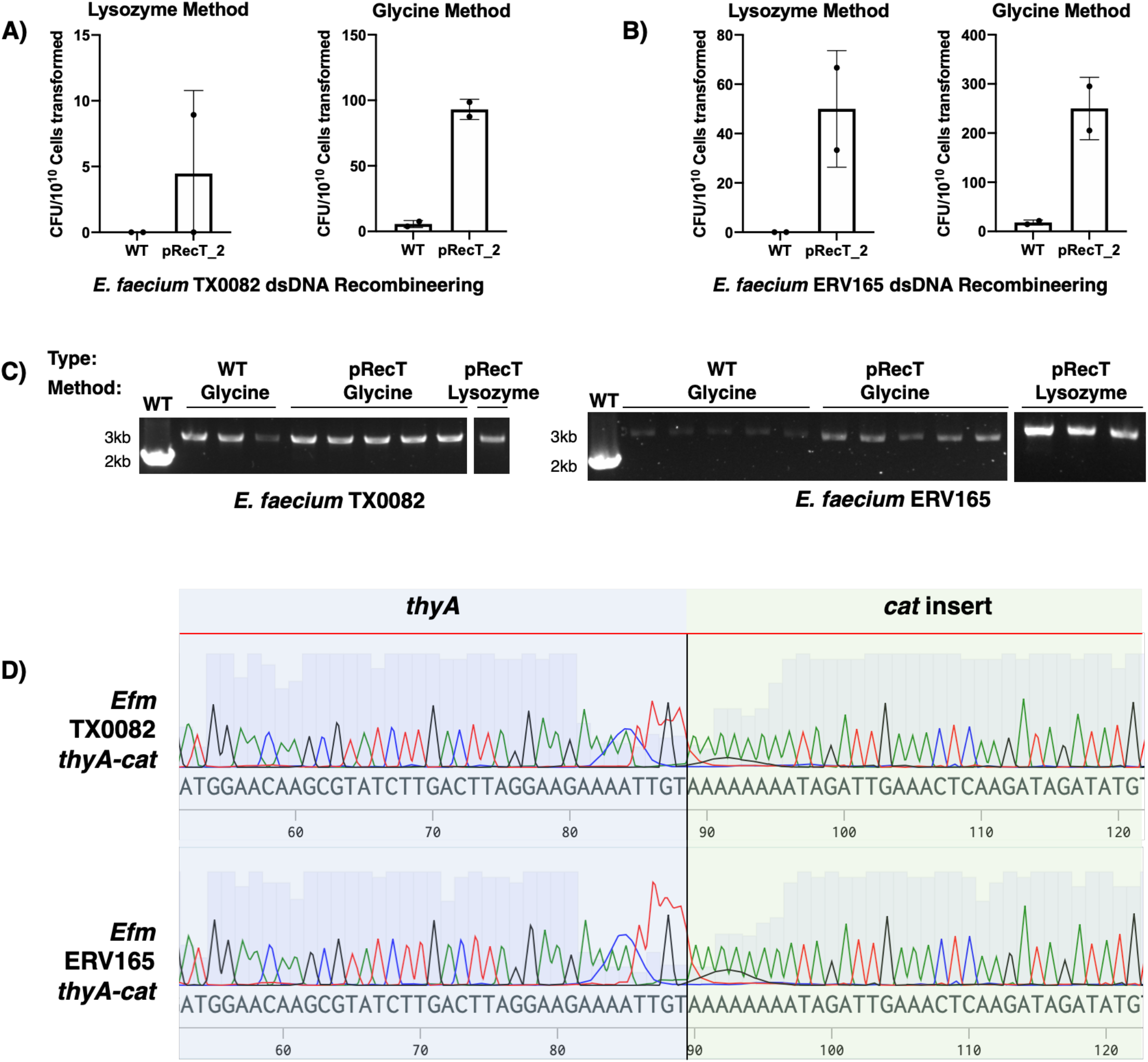
Recombineering technologies can be applied to pathogenic, multi-drug resistant strains of *E. faecium*. A) Relative CFU resulting from dsDNA recombineering to insert *cat* into *thyA* for vancomycin-resistant *E. faecium* TX0082 using both the lysozyme and glycine transformation methods. WT cells were used as a control to pRecT_2 harboring cells. B) Relative CFU resulting from dsDNA recombineering to insert *cat* into *thyA* for vancomycin-resistant *E. faecium* ERV165 using both the lysozyme and glycine transformation methods. WT cells were used as a control to pRecT_2 harboring cells. C) Representative DNA gel of colony PCR from resulting *E. faecium* TX0082 and *E. faecium* ERV165 colonies produced via dsDNA recombineering using the two transformation methods. PCR of WT *E. faecium thyA* of their respective strains were included as a control. D) Sequence identity of mutants show *cat* insertion in *thyA* at the correct location for dsDNA recombineering in both recombineered *E. faecium* TX0082 and *E. faecium* ERV165.

We then asked whether CRISPR-Cas9 can be used to produce scar-less mutations in VRE as was described in *E. faecium* Com15, *E. durans* and *E. hirae* (**Fig. 2 and S2**). Therefore, we transformed the VRE strains with pCas9-*thyA* and a ssDNA template designed to produce a 61 bp deletion using both lysozyme and glycine transformation methods. We were able to produce transformants using both transformation methods in *E. faecium* ERV165, with the lysozyme transformation method slightly more efficient than the glycine transformation method (**Fig. S5A**). To check for the introduced deletions, we performed PCR analysis on the *thyA* gene and found gel shifts corresponding to the 61bp deletion in 89% of the colonies picked in the lysozyme transformation method and 75% of the colonies picked in the glycine transformation method (**Fig. S5B**). As expected, all PCR products with apparent deletions had the exact 61 bp deletion (**Fig. S5C**). Taken together, this data suggests our developed recombineering technologies can be applied to pathogenic Clade A1 *E. faecium*.

## Discussion

In this study we developed recombineering and CRISPR-Cas9 technologies to enhance the efficiency of genomic engineering in both commensal and pathogenic *E. faecium*. Using these approaches, we were able to generate substitutions, deletions and insertions at high efficiency and fidelity on a relatively short timescale. We also demonstrated the utility of dCas9 for gene repression in *E. faecium*, thereby opening another method for investigating genes in *E. faecium*. By utilizing Cas9 selection, we generated scar-less substitutions and deletions using ssDNA oligonucleotides that can be readily purchased from commercial vendors (**Fig 1**). Thus, our method enables the functional assessment of any gene or protein in the *E. faecium* genome via genetic knockout, site specific mutations or whole protein domain deletions. The pRecT plasmid is readily lost from modified bacteria without erythromycin selection, and pCas9 can be cured upon cell passaging, allowing for sequential gene editing as well as the ability to isolate true isogenic mutants. This technology can be readily transferred to other *Enterococcus faecium* strains and *Enterococcus* species, allowing for other enterococci to be more genetically tractable. Moreover, the reagents used in this approach are highly modular, allowing for the replacement of *E. faecium* RecT with *recT* genes isolated from other species or the replacement of the plasmid backbone to further improve recombineering in other *Enterococcus* species and strains.

We discovered that *E. faecium* RecT can enable DNA insertions using dsDNA templates (**Fig. 4 and 6**). The successful insertion of *cat* allowed for the rapid screening of colonies, yielding multiple clones of high fidelity and on-target gene disruption. Studies are ongoing to further improve the capabilities of insertional mutation by optimizing CRISPR-Cas9 selection for shorter sequences that do not include antibiotic resistance. Nevertheless, the insertion mutation method provides a straightforward, facile means to produce gene knock-ins without the need of laborious passaging and screening. As technologies for custom long DNA synthesis improve and become more affordable, this technique will be powerful in providing a method to generate mutants in enterococci in a quick and easy fashion without the need for extensive cloning.

Although some methods to genetically engineer enterococci have been previously described^11,12,17^, our approach allows for mutant generation in *E. faecium* more efficiently and on a much faster time scale. Oligonucleotides can be ordered in advanced for both gRNA cloning and eventual ssDNA template delivery. Once available, gRNA can be constructed and cloned into pCas9 (day 1) and transformed into *E. coli* for propagation (day 2). Clones can then be analyzed for correct gRNA insertion (day 3) and grown for plasmid harvesting (day 4). Concurrently, pRecT harboring *E. faecium* can be prepared for recombineering by co-transforming harvested plasmid and ssDNA oligonucleotide template (day 4). Upon *E. faecium* colony outgrowth (days 5-6), clones can be screened by colony PCR and sequencing for mutant identification (day 7). True isogenic strains cured of CRISPR-Cas9/recombineering plasmids can then be generated by plasmid curing (days 8-9). As this process can be performed in parallel to analyze multiple targets, our recombineering method provides an efficient and potentially high-throughput means to characterize genes in *Enterococcus*.

## Acknowledgments

We thank Andrew Varble and Luciano Marraffini at The Rockefeller University for helpful suggestions and sharing plasmids. We thank Karthik Hullahalli and Kelli Palmer at University of Texas at Dallas for *Enterococcus* transformation tips and sharing plasmids. We thank Gregory Canfield and Breck Duerkop at University of Colorado Anschutz Medical Campus for *Enterococcus* transformation tips. We thank Katarzyna Cialowicz at the Rockefeller University Bio-imaging Resource Center for assistance with microscopy.

V.C. acknowledges support from The Rockefeller University Graduate Program and the National Institutes of Health (T32 A1070084). M.E.G thanks Hope Funds for Cancer postdoctoral fellowship and Melanoma Research Foundation for additional support. H.C.H. acknowledges support from the National Institutes of Health (NIGMS R01 GM103593 and NCI R01 CA245292) and Kenneth Rainin Foundation Synergy Award.

## Materials and methods

### Bacterial strains and growth conditions

*E. coli* DH5a strains were grown at 37°C in LB broth or LB agar. For liquid cultures, cells were grown in broth in shaking conditions at 220 RPM. Transformations were done according to manufacturer’s (New England Biolabs) instructions. Antibiotics for *E. coli* were used at the following concentrations: chloramphenicol, 10 μg/mL; erythromycin, 150 μg/mL; spectinomycin, 50 μg/mL. *Enterococcus* strains were grown at 37°C in Brain-Heart Infusion (BHI) broth or BHI agar. Antibiotics for *Enterococcus* strains were used at the following concentrations: chloramphenicol, 10 μg/mL; erythromycin, 50 μg/mL; spectinomycin, 250 μg/mL. For thymidine auxotrophy assays, *Enterococcus* strains were grown on MM9YEG agar (1x M9 salts, 0.25% yeast extract, 0.5% glucose, and 1.5% agar) supplemented with chloramphenicol and 40 μg/mL thymidine when appropriate. 10 μg/mL erythromycin was used for *Staphylococcus aureus* when propagating pRecT. *Staphylococcus aureus* was grown at 37°C in BHI broth or BHI agar.

### Bacterial transformations

*E. coli* transformations were performed using NEB 5-alpha chemically competent cells (New England BioLabs). *E. coli* transformations were performed according to manufacturer’s instructions. Transformations and plasmid extraction for *Staphylococcus aureus* was performed using a protocol described from a previous study^35^. *Enterococcus* transformations were performed via electroporation modified from a previously published methods^34,36,37^.

For the lysozyme transformation method, overnight cultures of *Enterococcus* were subcultured 1:100 in 25-50 mL of fresh media. Subcultures were grown until OD_600_0.6-0.8, then harvest by centrifugation 5000 RCF for 10 minutes at 4°C in a falcon tube. Supernatant was decanted, and the cell pellet was then resuspended in 1mL of ice cold 10% glycerol and transferred to an Eppendorf tube. Cells were pelleted by centrifugation at 7000 RCF for 8 minutes at 4°C. The supernatant was then aspirated, and the cell pellet was resuspended in 500 μL lysozyme mixture (10 mM Tris pH 8.0, 20% sucrose, 10 mM EDTA, 50 mM NaCl, and 30 μg/mL lysozyme) and incubated at 37°C for 20-30 minutes. Cells were pelleted again at 7000 RCF for 8 minutes at 4°C before aspirating the supernatant. Cell pellet was then resuspended in 1mL electroporation solution (0.5M sucrose and 10% glycerol). This step was repeated 3-4 times and resuspended in electroporation solution to the amount desired enough to complete the transformation. 100 μL aliquots of cells in electroporation solution was taken for each transformation. Remaining cells were stocked at -80°C to be used at a later date. Cells were transformed in 0.2 cm gap electroporation cuvettes (Bio-Rad) at 25 μF, 400 ohm and 2.5 kV. After electroporation, 400 μL SBHI, containing 0.5 M sucrose in BHI medium, was immediately added to transformed cells and left to recover for 3 hours at 37°C without shaking. Recovered cells were then plated on selective BHI agar plates.

For the glycine transformation method, overnight cultures of *E. faecium* were subcultured 1:50 in fresh 50 mL BHI broth containing 2% glycine and 0.5M sucrose again grown overnight. The next day, the cultures were pelleted again by centrifugation at 5000 RCF for 10 minutes, resuspended with 25 mL of prewarmed (37°C) BHI supplemented with 2% glycine and 0.5M sucrose, and incubated shaking at 220 RPM for 1-1.5 hours at 37°C. Cells were then pelleted again by centrifugation at 5000 RCF for 10 minutes, resuspended in 1mL of electroporation solution, and transferred to an Eppendorf tube. Cells were then pelleted again at 7000 RCF for 8 minutes at 4°C. The supernatant was aspirated, and the pellet was resuspended with 1 mL electroporation solution. This step was repeated once more before resuspending the cells in 1.2 mL of electroporation solution for transformation. 100 μL aliquots of cells in electroporation solution was taken for each transformation and kept on ice prior to electroporation. Remaining cells were stocked at -80°C to be used at a later date. Electroporations were performed in 0.2 cm gap electroporation cuvettes (Bio-Rad) at 25 μF, 400 ohm and 2.5 kV. After electroporation, 1 mL SBHI, containing 0.5 M sucrose in BHI medium, was immediately added to transformed cells and left to recover for at least 2 hours at 37°C without shaking. Recovered cells were then plated on selective BHI agar plates.

### Recombineering

Recombineering experiments were performed with desired *Enterococcus* strains transformed with pRecT or pRecT_2, plasmids containing an IPTG inducible RecT recombinase. The backbone for pRecT originates from a Gram-positive bacterial vector, pPM145, that is propagated from *Staphylococcus aureus* RN4220. The backbone for pRecT_2 originates from plZ12 propagated from *E. coli*. pRecT or pRecT_2 harboring cells were used to perform recombineering experiments using the electroporation protocols described above, with certain modifications. Specifically, for cells transformed by the lysozyme method, RecT was induced at 1mM IPTG when cultures reached an OD_600_ 0.3-0.4. The cultures were then returned to the incubator until they reached an OD_600_ of 0.6-0.8, and the protocol was continued as described above. For cells transformed by the glycine transformation method, overnight cultures grown in BHI with glycine and sucrose were pelleted and resuspended in fresh BHI containing 2% glycine, 0.5M sucrose, and 1mM IPTG. Cells were then incubated at 37°C for 1.5 hours for RecT induction before continuing with the wash steps in electroporation solution.

Recombineering-competent cells were transformed with either ssDNA template and accompanying pCas9 containing an appropriate gRNA targeting the desired mutation site, or dsDNA template. For ssDNA substitution recombineering, 100 base ssDNA template was designed with 49 base homology arms flanking a two-base substitution. For ssDNA deletion recombineering, 90 or 100 base ssDNA was designed with 45-50 base homology arms flanking the sequence to be deleted. All oligonucleotides used for transformation in this study were ordered from Integrated DNA Technologies and resuspended to a final concentration of 1 mM in MilliQ water. dsDNA templates were assembled by cloning the desired sequence into pET21 and amplified by PCR. PCR was performed using Q5 High-Fidelity DNA Polymerase (New England BioLabs) or PrimeSTAR Max DNA Polymerase (Takara). Prior to transformation of ssDNA templates, templates were dialyzed on MilliQ water using 0.025 μM membrane filter from Millipore (VSWP01300) for 15 minutes. Approximately 10-20 μL of dialyzed DNA template was used for each transformation. Typically, 3-4 μg of dsDNA were used for each transformation where applicable. After cell recovery, transformed cells were plated on BHI agar containing chloramphenicol. Resulting colonies were cultured, then screened for the appropriate mutations by PCR. For the PCR screening, 10 μL of culture were mixed with to 90 μL of TE buffer and heated to 95°C for 10 minutes. The cell debris was then pelleted by centrifugation at 20,000 RCF at 4°C for 5 minutes. 1 μL of supernatant was added directly to a PCR reaction.

### Plasmid construction

Cloning of plasmid constructs was performed using NEBuilder® HiFi DNA Assembly Master Mix from New England BioLabs. pCas9 was constructed using pVPL3004 as a backbone, replacing its encoded erythromycin resistance cassette with the chloramphenicol resistance gene, *cat*. Additionally, pUC19 origin of replication was inserted into the final pCas9 plasmid. pRecT was constructed using pPM145 as a backbone, inserting *recT* after an IPTG inducible promoter. pRecT_2 was constructed using plZ12 as a backbone, inserting *lacI* and the IPTG inducible *recT* from pRecT. Inserts were sequence verified via sanger sequencing through Genewiz. psrtA1 and psrtA2 were constructed using pKH12 as a backbone.

### Plasmid curing

To create isogenic mutants of *E. faecium*, pRecT and pCas9 were removed from the cells. This was accomplished by growing overnight cultures of newly generated mutants in antibiotic-free BHI, then subculturing the overnight cultures at 1:50 in fresh antibiotic-free BHI, incubating for several hours until cultures reach at least OD_600_0.8. Cultures were streaked on regular BHI plates for single colonies. Singles colonies were then picked and screened for antibiotic sensitivity by inoculating the colonies into liquid BHI with or without supplemented antibiotic. Resulting cultures containing antibiotic were then checked for lack of growth, indicating the plasmid from the original colony has been cured. The corresponding culture grown in regular BHI was then stocked for further experiments.

### gRNA cloning

gRNA cloning into pCas9 was accomplished by ligation of an annealed oligonucleotide pair into a BsaI-HFv2-digested pCas9. BsaI-HFv2 and T4 ligase were purchased from New England BioLabs. Generally, 4-5μg of pCas9 were digested with 100 units of BsaI-HFv2 overnight in a 50 μL reaction mix at 37°C. Oligonucleotide pairs containing the 30 base gRNA sequences with appropriate overhangs were ordered from Integrated DNA Technologies. To facilitate gRNA design, a script was written in Python to process DNA sequences (https://github.com/victorrchen/CRISPR). Oligonucleotide pairs were resuspended to 100 μM and annealed in a polynucleotide kinase (PNK) mix (New England BioLabs) consisting of 1.5 μL of each oligonucleotide, 41 μL of MilliQ water, 5 μL of PNK buffer and 1 μL of T4 PNK. This mixture was incubated at 37°C for 30 minutes, then supplemented with 0.5 μL of 5 M CaCl_2_ and transferred to a 95°C heat block for 5 minutes. The heat block was then removed from the heat source and allowed to cool at room temperature over the course of 3 hours. The oligonucleotide mixture was then diluted 1:10 to be used in the ligation reaction. For ligation, 20uL reactions were prepared with ∼500 ng of BsaI-HFv2 digested pCas9, 2 μL of diluted annealed oligonucleotide pairs, 2 μL DNA ligase buffer, 1 μL ATP, and 2 μL of T4 DNA Ligase in a 20 μL reaction mix. The reactions were incubated at 16°C overnight and transferred to 37°C for an hour before being heated to 80°C for 20 minutes. Ligated pCas9 was transformed into *E. coli* for plasmid propagation.

### Cas9 Western Blot

*Enterococcus* strains harboring pCas9 were grown overnight in 5mL BHI. Cells were pelleted at 5000 RCF and resuspended in lysis buffer (10% sodium dodecyl sulfate, 1x PBS, and 1x cOmplete^™^ Protease Inhibitor Cocktail in MilliQ water). Approximately 100 μL 0.1 mm diameter zirconium beads and 180 μL of 1x Laemmli Sample Buffer (Bio-Rad) were added to 2 mL screw cap micro tubes (Sarstedt Inc). 60 μL of resuspended cells were added to the tubes and bead beating was performed at max speed for 20 seconds in FastPrep FP120 cell disruptor. Beating was repeated twice. Tubes were then heated to 95°C for 10 minutes and spun down at 20000 RCF at 4°C for 5 minutes. 20 μL of supernatant was run on Stain-free^™^ gel (Bio-Rad). Gel was transferred onto nitrocellulose membrane. Membrane was blocked for an hour in block buffer (5% nonfat milk, 1x Tris-buffered saline, and 0.1% Tween 20). Immunostaining was performed using 1x HRP conjugated anti-Cas9 (Cell Signaling Technology 7A9-3A3) in block buffer overnight at 4°C on a rocker. Membrane was washed 3 times with TBS-T (1x Tris-buffered saline, and 0.1% Tween 20). Protein detection was performed using Clarity and Clarity Max ECL Western Blotting Substrates (Bio-Rad) on a Bio-Rad ChemiDoc Imaging System.

### Microscopy and Analysis

Microscopy and image quantification was performed as previously described^30^. Light microscopy was performed at the Rockefeller University Bio-imaging Resouce Center using Zeiss Axioplan 2 upright microscope on objective plan apochromat 63x oil NA 1.4 0.19mm. Images were captured using the equipped Hamamatsu High Resolution Digital B/W CCD Camera and acquired using MetaVue version 7.7.0.0. Images were processed using ImageJ. Bacterial cells to be imaged were grown overnight, pelleted, and resuspended in PBS, then were onto Fisherbrand Superfrost Plus Microscope Slides and covered with Zeiss cover glasses. Slides were sealed using Sally Hansen Insta-Dri Nail Color. Chain length was quantified live under the microscope and manually counted. Approximately 300 particles were picked for each condition, where one particle is defined by one continuous chain of bacteria. Statistical analysis was performed on PRISM 9 using the Kruskal-Wallis test with Dunn’s correction.

