## Supplemental_figures for "RecT recombinase expression enables efficient gene editing in *Enterococcus*"

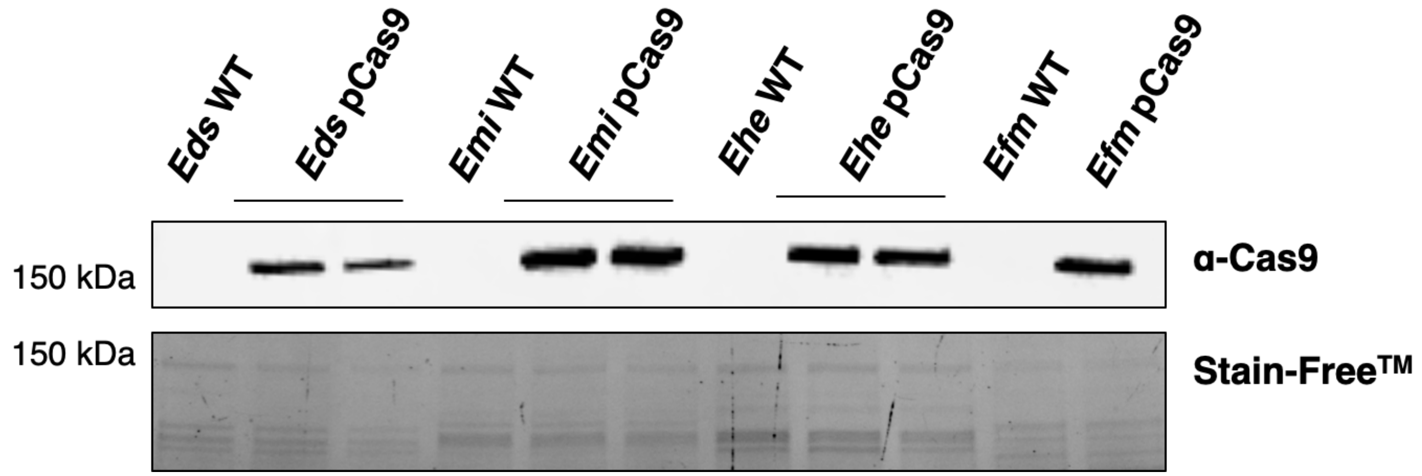


**Figure S1**. **Western blot for Cas9 in *Enterococcus*.** *Enterococcus* species were transformed with pCas9 and assayed for Cas9 expression. From left to right, *E. durans* (*Eds*), *E. mundtii* (*Emi*), *E. hirae* (*Ehe*) and *E. faecium* Com15 (*Efm*) are shown.

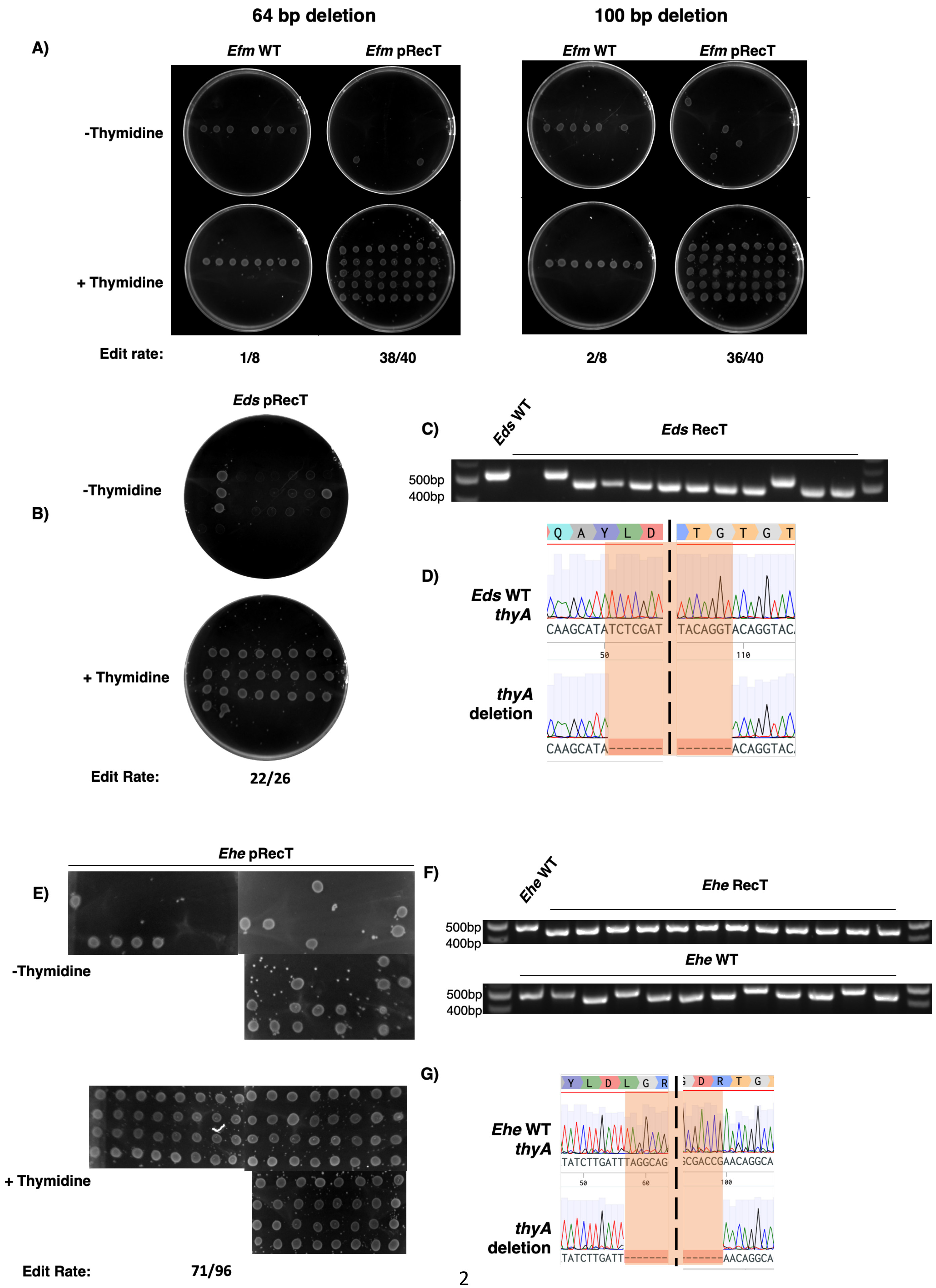

64 bp deletion
100 bp deletion
A)
Efm WT
Efm pRecT
Efm WT
Efm pRecT
-Thymidine
+ Thymidine
Edit rate:
1/8
38/40
2/8
36/40
Eds pRecT
B)
-Thymidine
+ Thymidine
Edit Rate:
22/26
C)
Eds WT
Eds RecT
500bp
400bp
D)
Q A Y L D
T G T G T
Eds WT thyA
CAAGCATATCTCGAT
TACAGGTACAGGTAC
50
110
thyA deletion
CAAGCATA--------
-------ACAGGTAC
Ehe pRecT
E)
-Thymidine
+ Thymidine
Edit Rate:
71/96
F)
Ehe WT
Ehe RecT
500bp
400bp
Ehe WT
500bp
400bp
G)
Y L D L G R
D R T G
Ehe WT thyA
TATCTTGATTTAGGCAG
CGACCGAACAGGCA
50
60
100
thyA deletion
TATCTTGATT------
------AACAGGCA

**Figure S2. Spot assays for *E. faecium* and analyses for other *Enterococci* mutants.** A) Spot assay was performed for *E. faecium* exconjugants resulting from CRISPR-Cas9 mediated recombineering of *thyA* deletion mutants. *E. faecium* was spotted on MM9YEG agar with chloramphenicol, with or without thymidine. B) Spot assay for *E. durans* exconjugants from CRISPR-Cas9 mediated recombineering to delete *thyA*, performed as previously described. C) DNA gel of *E. durans* mutants compared to WT. D) Sequence identity of *thyA* mutants from *E. durans*. Expected deletion size of 58bp is shown. E) Spot assay for *E. hirae* exconjugants from CRISPR-Cas9 mediated recombineering to delete *thyA*, performed as previously described. F) DNA gel of *E. hirae* mutants compared to WT. G) Sequence identity of *thyA* mutants from *E. hirae*. Expected deletion size of 58bp is shown.

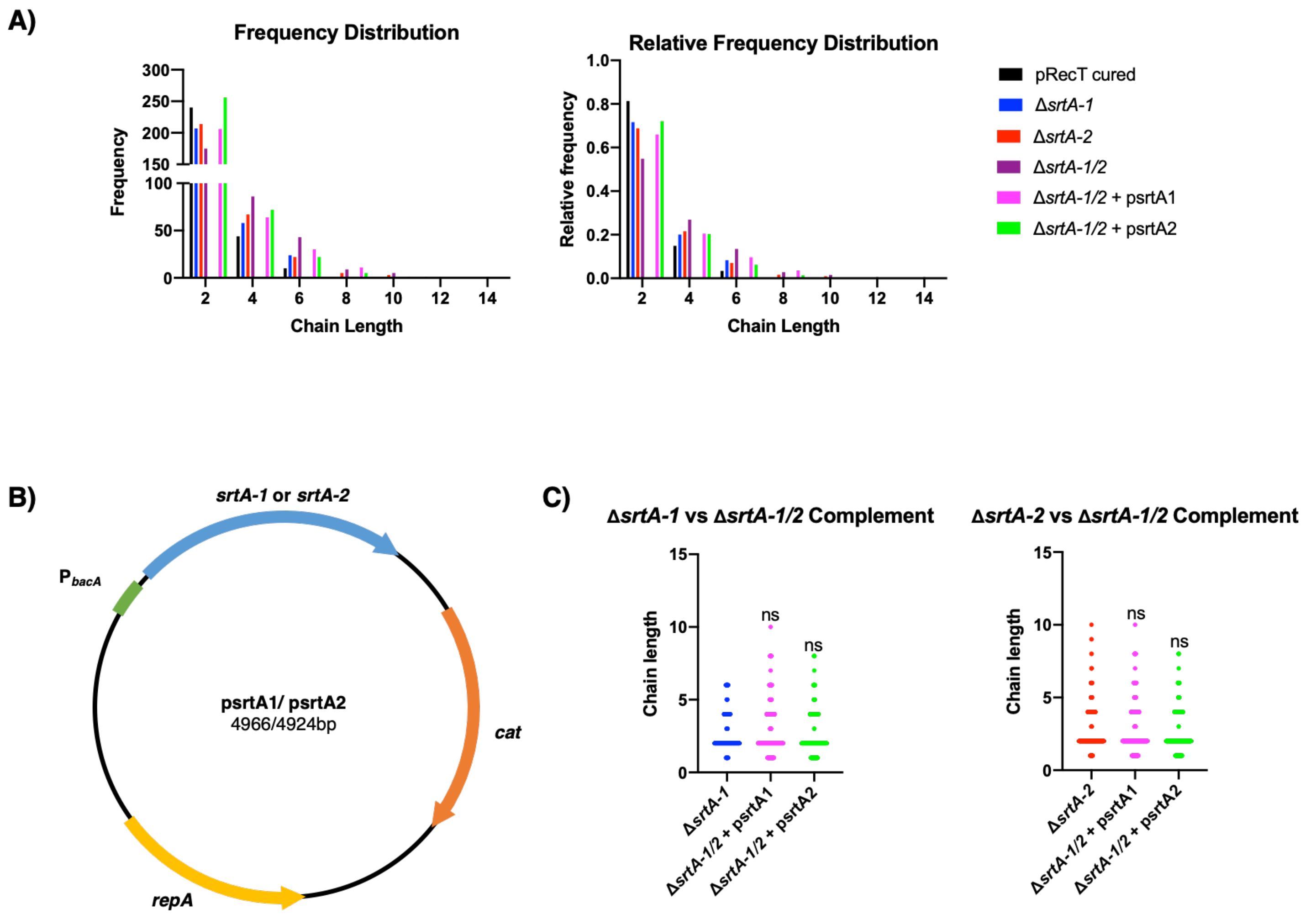


**Figure S3. Extended analysis of sortase A related cell chaining in *E. feacium* Com15.** A) Absolute and relative frequencies of chain lengths observed in control, sortase A mutants and sortase A complement strains were graphed using a bin width of two. Relative frequencies were calculated by taking the absolute frequencies and dividing it by the total number particles picked for each condition. B) Plasmid map depicting the complementation plasmid used. Plasmid backbone was taken from pKH12. *srtA-1* or *srtA-2* were cloned and put under the control of a constitutive promoter, $P_{bacA}$. C) Extended graphs of analyses on cell chaining from Kruskal-Wallis ANOVA with Dunn's correction corresponding to Figure 5E. Comparisons for *E. faecium* Δ*srtA-1* and Δ*srtA-2* to Δ*srtA-1/2* strains with complementation plasmids are shown.

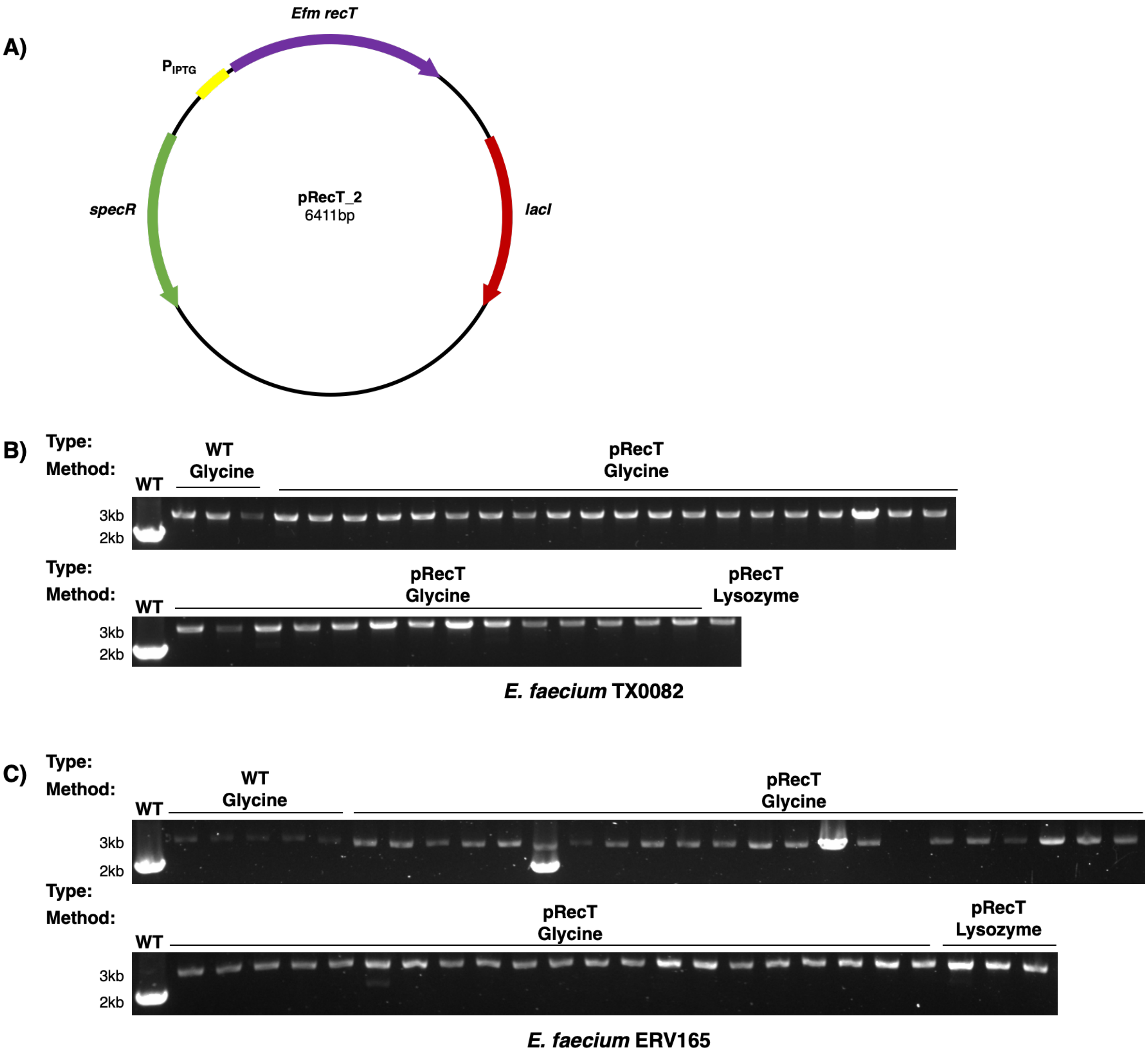


**Figure S4. Recombineering can be applied to vancomycin-resistant *E. faecium*.** A) Plasmid map of pRecT_2. *recT* operon from pRecT was transferred to pIZ12 backbone to create pRecT_2. B) Uncropped DNA gel of colony PCR from resulting colonies produced in *E. faecium* TX0082 dsDNA recombineering using the two transformation methods. Gel is a continuation from Figure 5C. C) Uncropped DNA gel of colony PCR from resulting colonies produced in *E. faecium* ERV165 dsDNA recombineering using the two transformation methods. Gel is a continuation from Figure 5C.

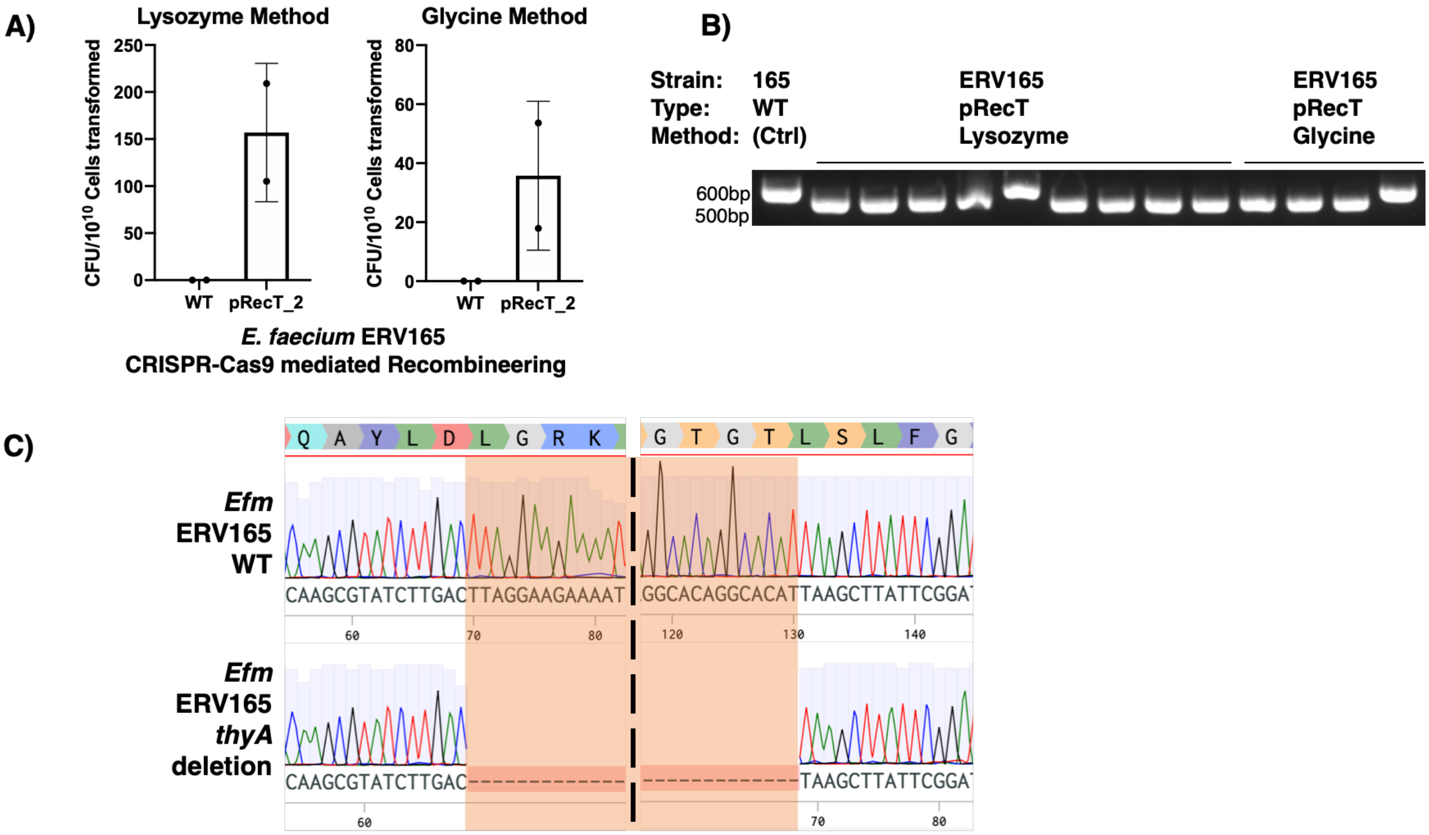


**Figure S5.** ***E. faecium*** **ERV165 CRISPR-Cas9 mediated recombineering.** A) Relative CFU resulting from CRISPR-Cas9 mediated recombineering to produce short deletions in *thyA* for vancomycin-resistant *E. faecium* ERV165 using both the lysozyme and glycine transformation methods. B) DNA gel of colony PCR of resulting *E. faecium* ERV165 colonies via CRISPR-Cas9 mediated recombineering using the two transformation methods show 61bp deletions in *thyA* for most of the colonies picked. C) Sequence identity of *thyA* deletion mutants in *E. faecium* ERV165 to be of high fidelity, with exactly 61bp deleted.

| Chain Length Bins | pRecT cured | Δ*srtA-1* | Δ*srtA-2* | Δ*srtA-1/2* | Δ*srtA-1/2* + psrtA1 | Δ*srtA-1/2* + psrtA2 |
|---|---|---|---|---|---|---|
| 1 | 35 | 17 | 21 | 20 | 43 | 55 |
| 2 | 205 | 190 | 193 | 155 | 163 | 201 |
| 3 | 10 | 8 | 5 | 2 | 9 | 2 |
| 4 | 34 | 50 | 62 | 84 | 55 | 70 |
| 5 | 4 | 5 | 11 | 16 | 12 | 9 |
| 6 | 6 | 19 | 11 | 27 | 18 | 13 |
| 7 | 0 | 0 | 3 | 0 | 1 | 1 |
| 8 | 0 | 0 | 2 | 9 | 10 | 4 |
| 9 | 0 | 0 | 2 | 0 | 0 | 0 |
| 10 | 0 | 0 | 1 | 5 | 1 | 0 |
| 11 | 0 | 0 | 0 | 0 | 0 | 0 |
| 12 | 1 | 0 | 0 | 0 | 0 | 0 |
| 13 | 0 | 0 | 0 | 0 | 0 | 0 |
| 14 | 0 | 0 | 0 | 1 | 0 | 0 |

**Table S1. Raw data of chain length frequencies of different strains of *E. faecium* Com15.** Table depicts raw data used to produce graphs in Figure 5E, S3A and S3C. Numbers represented the number of particles counted corresponding to the chain length between 1-14.
